# Climate or disturbance: temperate forest structural change and carbon sink potential

**DOI:** 10.1101/478693

**Authors:** Travis Andrewsa, Michael Dietzeb, Robert Boothc

## Abstract

Anticipating forest responses to changing climate and disturbance regimes requires understanding long-term successional processes and aggregating these local processes into global relevance. Estimates of existing forest structure and biomass are improving globally; however, vegetation models continue to show substantial spread in predictions of future land carbon uptake and the roles of forest structural change and demography are increasingly being recognized as important. To identify mechanisms that drive change in tree size, density, and carbon, we need a better understanding of forest structural trajectories and the factors that affect those trajectories. Here we reveal a coherent, cyclic pattern of structural change in temperate forests, as predicted by successional theory, and identify significant sensitivity to climatic precipitation and temperature anomalies using large datasets and empirical modeling. For example, in the eastern US above average temperature (+1°C) was associated with a 27% (−0.4±0.1 Mg C ha^-1^ yr^-1^) reduction in productivity attributed to higher rates of disease (+23%), weather disturbance (+57%), and sapling mortality. Projections of future vegetative carbon sink potential suggests biomass would be lowest on managed lands (72±2 Mg C ha^-1^) and highest when larger trees survive in undisturbed conditions (153±21 Mg C ha^-1^). Overall, the indirect effects of disturbance and mortality were considerably larger than the direct effects of climate on productivity when predicting future vegetative carbon sinks. Results provide robust comparisons for global vegetation models, and valuable projections for management and carbon mitigation efforts.

## INTRODUCTION

Regional-scale shifts in forest structural characteristics have long-term implications for global terrestrial carbon stocks, but remain a challenge to predict due to the complex interactions of succession, disturbance, and climate sensitivity. Ecological succession is one of the most enduring concepts of forest ecology, describing how resources, competition, and disturbance influence tree establishment, growth, and mortality. As succession proceeds, resource limitation, climate, and disturbance bound the possible combinations of tree size and stem density, though these relative influences remains poorly constrained (Seidl *et al.*, 2011). Given this uncertainty, even anticipating relatively simple climate-induced structural changes in forests, such as changes in stem density or carbon accumulation, are challenging using traditional ecological approaches, which have often been spatially and temporally limited (Foster *et al.*, 2010). Furthermore, spatial heterogeneity in land use has led to considerable variability in forest ages (Caspersen *et al.*, 2000, Pan *et al.*, 2011). Climate change will play out against this variable mosaic, which for much of the world is not at a landscape-scale structural steady-state (Turner *et al.*, 1993).

Of the many factors that influence forest structure and carbon accumulation, the response of forests to changing climate is of particular concern (Anderegg *et al.*, 2013, Change, 2014, Dale *et al.*, 2016, Pan *et al.*, 2011, Purves & Pacala, 2008, Schellnhuber *et al.*, 2012, Seidl *et al.*, 2011). For example, the direct effect of warming is generally to increase plant productivity, but indirectly higher temperatures are widely associated with drought stress, increased pathogen and insect infestations, increased tree mortality and selective recruitment (Anderegg *et al.*, 2013). A challenge for climate mitigation planning and forest management is to predict future carbon sequestration and density of forests recovering from historic land use in the context of climate change (Caspersen *et al.*, 2000) and changing disturbance regimes. Uncertainty about changing climate impacts on temperate forest steady state conditions is a source of risk to sustainability goals and carbon finance. Temperature, precipitation, disturbances, and succession all have non-linear effects that are not reliably characterized at regional to continental scales by small-scale experiments or current process-based models. Here we empirically quantify successional structural processes and identify the influence of changing climate and disturbances on tree growth, density, and carbon accumulation. The findings provide a fundamental understanding of ecological processes that can benchmark vegetation models and help inform forest carbon investment.

## METHODS

To quantify structural change we use the US Forest Service Forest (USFS) Forest Inventory and Analysis (FIA) database to map (in phase space) how stem density, radial tree growth, and productivity (change in above and below ground vegetation carbon) change as a function of mean diameter and relative density (i.e. stocking; Arner et al., 2001) over time for a broad range of common temperate forest densities. These relationships form the empirical succession mapping (ESM, Fig. 1). The ESM builds on classic forestry research that used mean diameter and stem density to compare the growth and expected structural trajectory of even-aged single-species forests (Reineke, 1933, YODA, 1963). The ESM was used as the basis for comparative and predictive models to assess the influence of climate and disturbance on forest structure and carbon accumulation. A schematic overview of methodology is provided in Table 1 that shows the ESM is a set of simple analyses rather than a complex model.

**Table 1:**
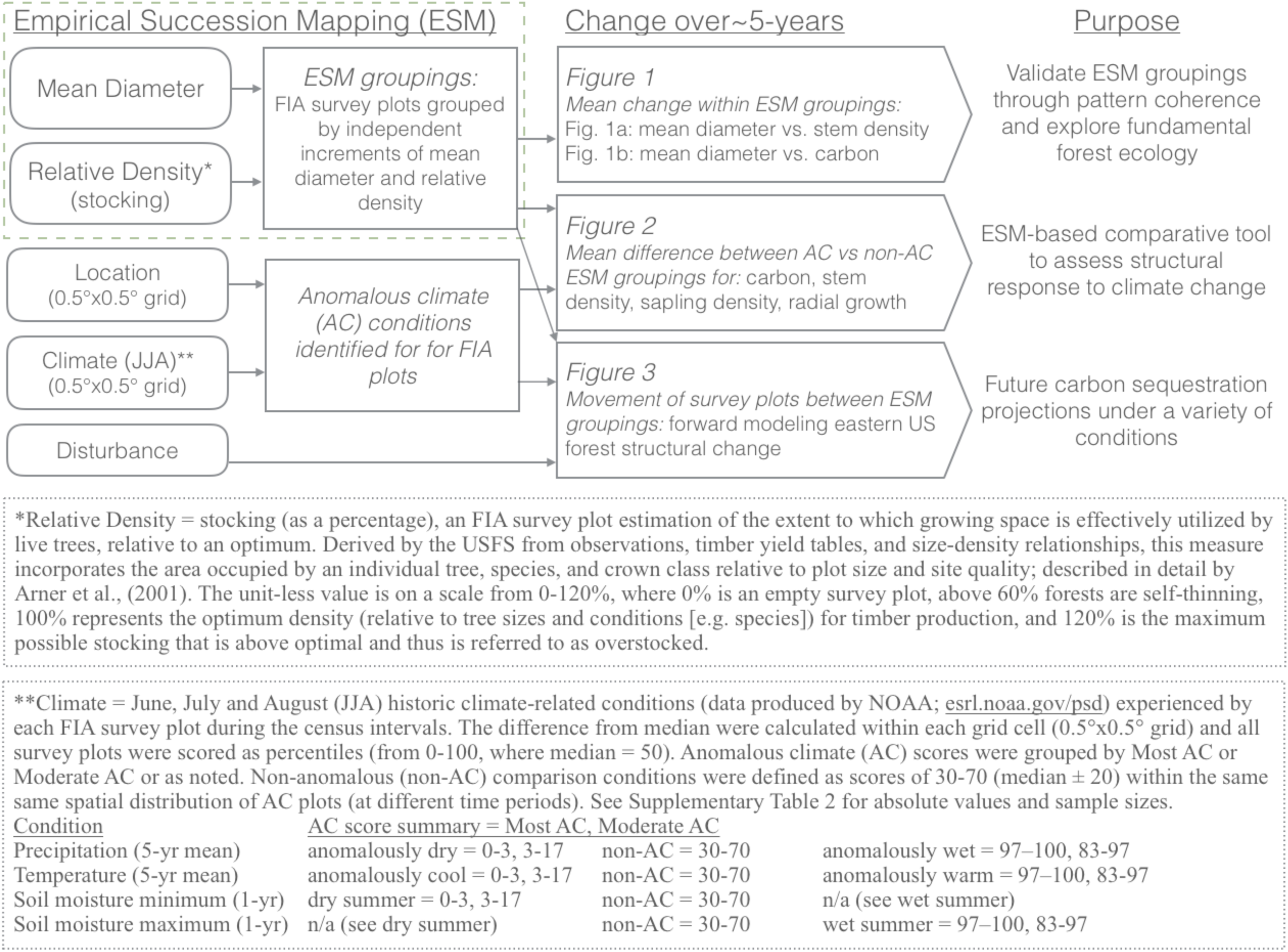
Methods Summary Schematic

**Figure 1.**
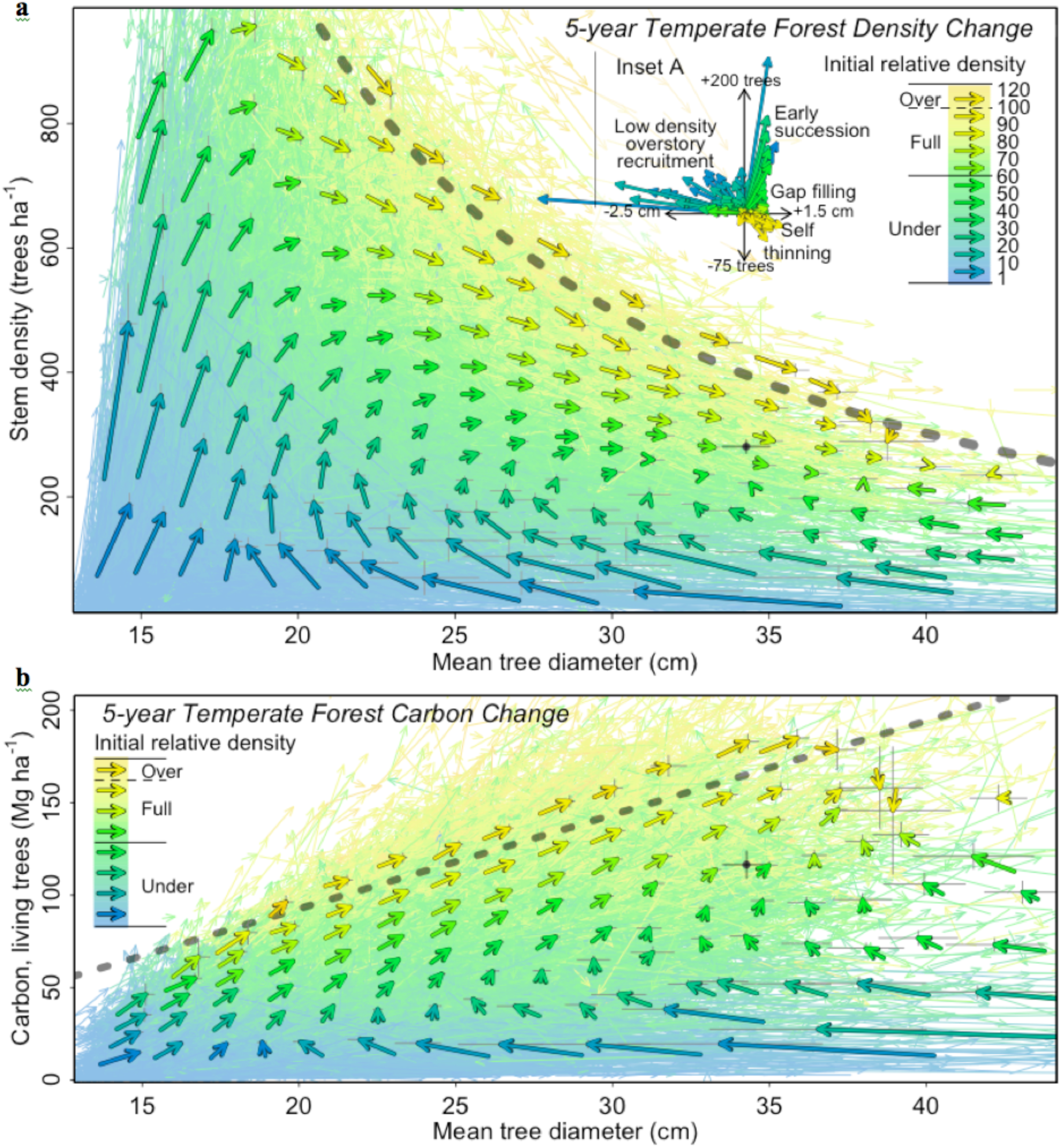
Empirical Succession Mapping for a) density and b) carbon. Measured changes in the number, size, and carbon stored in trees (n = 34,404) were independently averaged at increments of initial mean diameter and initial relative density (bold vectors) with bootstrapped error (±2SE; 1000 iterations, thin grey lines); except for forests with initial mean diameters greater than 32 cm, where moving averages were used due to high probability of mortality (30%) and recruitment (35%) making forests with these large mean diameters unlikely (4.5% of dataset). Inset A compiles mean density change vectors and highlights likely ecological processes driving observed changes. Grey points show projected mean steady-state density and carbon under current conditions with uncertainty. The best fit delineation between fully-stocked and over-stocked forests in a) was a power law in the form of Equation (S1), with a scaling exponent of b = -1.75, and in b) was linear (slope = 5.6, y-intercept = -19.4).

ESM development and analyses were based on repeated forest measurements and climate data as described below. All data filtering, statistical analyses, and figure creation were performed using R version 3.1.2 (Team, 2012). The annotated R code and filtered dataset used for all analyses are available on Github (github.com/wanderswest/ESM-FIA). All raw data is publicly available with links provided in the description below.

### DATA

#### USFS Forest Inventory and Analysis

The US Forest Service has measured or calculated many characteristics associated with millions of trees in a regular gridded sampling of forests across the US. Over the last decade these measurements have been standardized and repeated for forest plots on an approximate 5-year cycle. Measurements are made for trees larger than 12.7 cm diameter at breast height (DBH) on 0.067 ha plots (subplots). We subset the FIA (*http://apps.fs.fed.us/fiadb-downloads/datamart.html*) using the criteria in Supplementary Table 1. FIA surveys are performed at the state level and due to the high-quality standards, Louisiana, Florida, and West Virginia have less dense sampling than adjacent states. Sampling heterogeneity was not expected to impact analyses presented here that broadly examine relative forest change rather than attempting to characterize a specific species or location. Environmental characteristics (e.g. soil quality, microclimates) are expected to have a consistent influence on forest structure during the census interval.

We filtered the FIA to 38,857 forest interior plots composed of ~1.0 million trees (minimum diameter 12.7 cm) in the US east of -95W longitude, resurveyed at a 5.0±0.01 (mean±SE) year census interval between 1998-2012 using the same survey method. We excluded seedlings, planted forests, and forests with edge effects (see criteria in Supplementary Table 1). Saplings (2.54-12.7 cm diameter) were excluded except for carbon calculations and as a response variable to climate conditions. Plots with indications of management (e.g. harvesting, thinning) during the census interval (n = 4452) were used only in the “with logging” future carbon sink projection. Vegetation carbon was estimated in the FIA for above and below ground portions of living trees greater than 2.54 cm in diameter, excluding foliage based on allometric relationships (Jenkins *et al.*, 2003). Productivity was calculated for each plot as the difference between carbon stored in living biomass at the initial survey and second survey divided by the length of the census interval in years. Forest carbon content was considered a structural characteristic and thus productivity represents a measure of structural change. Soil carbon sequestration is an important component of the carbon balance in forests that was not assessed by this study.

Plot-scale disturbance was denoted in the FIA as AGENTCD, identified by technicians in the field. Here we included disturbances that caused mortality of one or more trees during the census interval associated with insects, disease, fire, animals, and/or weather. Specifically, disturbed trees were alive at the initial survey and dead at the resurvey with the AGENTCD identified as the cause of mortality, which impacted 31.0% of all forest plots and 34.3% of self thinning plots (i.e. relative density > 60%). Mortality associated with vegetative suppression/ competition and unknown causes were expected to be largely associated with autogenic succession and were not considered disturbance. Notably, unknown causes of mortality were common, attributed to 45.4% of tree mortalities. Since disturbances (fire, insect, disease, animal, weather) may be difficult to identify and attribute to mortality in the field, we expect disturbances were conservatively estimated. FIA surveys are run at the state level and local knowledge of hurricane impacts or extensive droughts, for example, would aid technicians in identifying weather related mortality.

Identification of tree mortality causes can be both difficult and subjective, and previous studies have identified the drawbacks and value of FIA determinations (Kromroy *et al.*, 2008, Meng *et al.,* Stewart *et al.*, 2009). Similar to previous studies, we identified and utilized relative change in disturbance frequencies. Overall, our main conclusions regarding the influence of climate on forest structure and carbon sink potential did not require high levels of disturbance-identification accuracy. Furthermore, we found that excluding all forests with unknown disturbances had minimal influence on frequencies of disturbances associated with climate anomalies (see Supplementary Table 3).

#### Climate conditions data

We used monthly 0.5° × 0.5° gridded datasets of historic climate data for 5-year mean precipitation (PREC/L) and surface temperature (GHCN_CAMS 2m), and 1-year maximum/minimum soil moisture (CPC), produced by NOAA from land-based instrument records spanning the last century (esrl.noaa.gov/psd). In the Eastern US, rolling 5-year average June, July, and August (JJA) summer climate conditions over the past two decades were identified for each grid cell. Summer climate was chosen as the most relevant period of growth and climate induced stress for forests of the Eastern US. For the one-year summer wet and dry anomalies, soil moisture was chosen as a more accurate measure than precipitation to infer forest water stress during extreme events. The soil moisture data is derived from a simple bucket model that integrates observations of temperature and precipitation to model water height equivalent in the top meter of soil. Soil moisture z-scores were calculated for each grid cell using the period from 1991-2012. Specific definitions of climate conditions used in each analysis are in Supplementary Table 2. Numerous additional climate condition comparisons, such as temperature variability and nighttime temperatures, could be assessed in future work.

Climate conditions during the census interval were assigned to each forest plot. Due to the dense and on-going nature of the FIA program, within each 0.5° × 0.5° grid cell, several forest plots have been surveyed over several different time periods. Here we exploit differences in climate between different survey periods within each grid cell to determine forest plots that experienced anomalous climate. See Table 1 and Supplemental Information for more details on how climate anomalies were calculated. Most AC (AC scores: 0-3 or 97 -100) and Moderate AC (AC scores: 3-17 or 83-97) were chosen to correspond to +0.5°C and +1.0°C warming, respectively; and, then used to make comparisons to disturbance and other climate conditions (Supplementary Tables 2, 3 & 4). As noted, other AC score groupings were utilized to highlight relationships. The future projections steady-state model combined Most AC, Moderate AC and slightly AC, or approximately AC scores of 0-40 or 60-100 (Supplementary Table 2), for the increased data density needed to cycle forest densities (Oliver & Larson, 1996).

### PHASE SPACE MAPPINGS

#### Empirical succession mapping

To quantify forest structural change we used the filtered FIA dataset to map how stem density, radial tree growth, and productivity change as a function of mean diameter and relative density over time. The mean plot-level data were averaged for comparison using independent groupings of mean diameter versus relative density at the initial census. Relative density is known in the forestry community as stocking. It was renamed here for broader understanding. Stocking is the FIA plot estimation of the extent to which growing space is effectively utilized by live trees, relative to an optimum (Arner *et al.*, 2001). Grouping forest plots by relative density at initial census, rather than stem density, allows for comparison across forest types. The relative comparison of tree volume across forest types nationwide is a core mission of the FIA. Methods of comparison were first widely adopted in the 1950’s and have since been revised to incorporate the area occupied by an individual tree, species, and crown class relative to plot size and site quality. The associated stocking algorithms, derived from observation-based forest studies, timber yield tables, and size-density relationships, are described further by (Arner *et al.*, 2001). Mean diameter is related to stand age, however it is expected to be a superior proxy for successional status (when used in combination with relative density), particularly in mixed age forests. Notably, the 500 forests believed to be the oldest, 122-255 years old, had a wide range of mean diameters, which suggests the steady-state density of an individual late successional forest follows a long-term, large-scale circulation of densities.

#### Averaging across different species and locations

Resource limitation influences all tree species to follow size-density relationships, although the specific maximum constraint on this relationship remains debated (Enquist & Niklas, 2001, Zeide, 2010). As one of the earliest quantified relationships in ecology, the -3/2 self thinning power law (YODA, 1963) describes an upper boundary of plant size-density relationships. Forests compensate for resource limitation through thinning such that mean diameter growth is offset by a constant proportional reduction in stem density, thus following a power law relationship (Equation 1) with a scaling exponent (b) of -1.5 or -2 for the self thinning rule or metabolic scaling (Enquist & Niklas, 2001, Zeide, 2010), respectively. Y_0_ is an ecosystem dependent constant. The scaling exponent describes the balance of recruitment, radial growth, and mortality on forest structural trajectory.

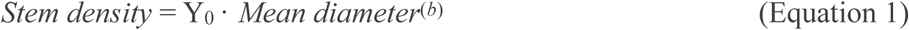

In the ESM we quantified structural change approaching maximum size-density constraints; and, despite a wide variety of temperate forest species and locations, a coherent pattern of independent structural change vectors emerged that indicated average change was robust and predictable at a regional scale. While individual forest plots had a wide range of structural responses (i.e. recruitment, mortality, and radial growth), when 10 or more plots were grouped by both mean diameter and relative density, there were strong structural trajectories across species and locations with low uncertainty. The key for accurate comparison was utilizing the relative density measure (i.e. stocking) that was developed by the USFS specifically for the purpose of comparing diverse forest types (Arner *et al.*, 2001).

As a comparative tool, ESM is useful to identify the influence of factors that are independent of species (e.g. anomalous climate) and comparable across similar locations, such that the distribution of species remains heterogeneous and the influence of geographic gradients (e.g. annual temperature) remains consistent (see Supplemental Information for further description of the analysis framework). The drawback is unique species specific responses are incorporated into error and results describe the net regional effect. In other words, the ESM predicts the regional response but not the exact response at specific locations or for individual species, which are well known to be variable.

#### The influence of climate anomalies

To assess the ESM sensitivity to climate, we examined structural change that occurred during anomalous precipitation, soil moisture and temperature conditions. Forest plots that experienced anomalous climate (AC) were compared, using spatially weighted ESM groupings (further described in Supplemental Information), to forests that experienced near median conditions (30^th^ -70^th^ percentile; non-AC conditions). AC conditions were not evenly distributed across the eastern US and thus the non-AC comparison plots were selected to represent the same geographic area as the AC plots. Because the ESM represents structural change across time, comparisons of different time periods should not influence the results. For each anomalous climate comparison, the non-AC ESM was also compared to a subset of itself to check that differences and error were minimal. Interactions between AC conditions and other AC conditions or disturbances were assessed for Moderate and Most AC conditions (see Tables S2 & S3).

#### Forward modeling

We used plot-level change to understand how the distribution of forest characteristics would change through time across ESM groupings. The ESM carbon mapping was used as a basis for projecting current forest change into the future using a data-driven approach and a space for time substitution. This forward model is merely a computationally intensive weighting process of existing empirical data and no synthetic plot-level data were manufactured. This method identified how plots changed ESM groupings from initial survey to resurvey (~5-years later). The distribution of plots across the ESM groupings at the resurvey was then used to select plots with the same distribution at the first survey. The process was then iterated with each iteration representing 5-years of changing distribution of forest structural characteristics. At each iteration the newly selected plots were randomly chosen within each ESM group based on the number of plots that entered the ESM grouping in the previous iteration. From this selection process, the average change in mean diameter, carbon and stem density were recorded. The ESM groupings were defined as independent increments of 0.5 cm mean diameter and 10 relative density; up to overstocked forests (relative density >100%), which were grouped together as the largest relative density increment. To avoid missed connections in forests with mean diameters larger than 32 cm, the ESM groupings were enlarged to increments of 2 cm mean diameter and 20 relative density; and all forests with mean diameters greater than 40 cm were put into the 40 cm mean diameter grouping. Carbon projections were not sensitive to a range of ESM grouping increment sizes tested. Within each ESM grouping, if more vectors ended in the ESM group than started, start plots were randomly repeated. Finally, in the rare occurrence that any vectors ended without trees, they were started in the ESM grouping with increments of the smallest mean diameters and lowest relative density. This process of restarting forest growth assumes the presence of large saplings, which if saplings are not present or if the land will not return to forest, may slightly overestimate or underestimate average eastern US forest density, respectively.

Each disturbance and climate scenario started with the “all forests” dataset (excluding logged forest plots, n = 34,405) that then was used to select survey plots of forests that experienced conditions specified by that scenario. Scenarios were created by excluding plots without those climate or disturbance conditions (i.e. the 2x disturbance condition dataset is the “all forests” dataset that randomly excluded 50% of the undisturbed forest plots). For consistency, the scenario with logging was also started using “all forests” and at the first iteration, plots with logging were added. Climate scenarios were ramped up in intensity during the first four model iterations (20-year linear ramp) to mimic slow projected change in climate over the next couple decades. Climate change projections were the maximum intensity that reached a steady-state density in projections, which were generally mild conditions because more extreme conditions (e.g. +1.0°C warming) did not have enough data to represent the natural cycling of forest density (Oliver & Larson, 1996) and therefore did not reach steady-state. Climate scenarios were considerably less extreme than those expected in the distant future but were useful to assess controls on steady-state density. The model was run for 60, 5-year iterations and most scenarios reached a density steady-state within 150 years. Model uncertainty was assessed by running each scenario 10 times using equal-sized datasets (n=7202; i.e. 95% of the smallest dataset) randomly selected at each iteration and then taking the standard deviation of the final steady-state densities (each based on the final 10 iterations).

#### How are species/locations/succession differences incorporated into future projections?

Eastern US forests are largely recovering from historic land use. Disturbances, mortality and recruitment of late succession species are also ongoing, which effectively cycles forest densities to produce a landscape-scale steady state (Turner *et al.*, 1993). Taking an average of all FIA plots, as is done at the start of the future projections (Fig. 3a), underestimates forest carbon sink potential due to the overabundance of plots recovering from land use change relative to those recovering from natural disturbances. Thus in simplest terms, the forward model intends to select forest plots that represent part of the natural density cycle (or with logging), from which it calculates the average mean diameter and carbon of the selected forests. In each scenario, the model found approximately a third of FIA plots (i.e. ~13,000 of the 34,405 starting plots in the “all forests” scenario) represent a steady-state cycling of density, however, at each iteration, plots are selected randomly within the ESM mean diameter/relative density increment groupings; and thus the full range of modern species/locations should be represented in the final steady-state equilibrium. Notably, plots with the same relative density may vary widely in growth rate depending on species and site quality. Connecting plots with rapid growth to those with slow growth (and vice versa) would be a poor model for an individual forest, however, when averaged across all plots, the average of all structural changes should remain an accurate representation of the average for the projected eastern US, even accommodating shifts in species over time; see Supplemental Information for further explanation. Data from each scenario (n >7500) was well distributed across the eastern US, and thus should represent comparable average forest structural change.

## RESULTS

The ESM reveals mean tree diameters and stem density change in a coherent, cyclic pattern predicted by successional theory (density mapping, Fig. 1a). The ESM uses the wide range of individual plot structural responses (i.e. recruitment, mortality, and radial growth) associated with the immense complexity of species/environment/disturbance conditions across eastern US forests and identifies average structural changes that are robust and predictable at this broad spatial scale (further explained in Methods). The ESM patterns generally highlight the primary processes likely driving change, including early successional establishment and growth, self-thinning, and gap filling (Fig. 1a inset).

The majority of forests in the eastern US are recovering from past disturbance and/or management, resulting in increased stem density and accumulated carbon during the census interval (+4.2±0.1 stems ha^-1^ yr^-1^, +1.7±0.01 Mg C ha^-1^yr^-1^; respectively). In Fig. 1a, vectors with the largest vertical component occur in forests with small mean diameters, which represents the early successional tree recruitment (+73.6±15.4 stems ha^-1^ yr^-1^; i.e. crossing the 12.7 cm diameter threshold) and maximum carbon accumulation in the mapping (+2.4±0.7 Mg C ha^-1^yr^-1^; Fig. 1b). By contrast, at larger mean diameters (i.e. > 18.5 cm) in forests with low stem densities (relative density < 20%), recruitment (+10.3±0.3 stems ha^-1^yr^-1^) resulted in sharp reductions of mean diameter over the census interval. These mean diameter reductions are analogous to the classic transition from mid to late successional structure when smaller young trees are recruited into low density stands with sparse large trees. These late successional forests would typically maintain lower relative densities. At higher stem densities (i.e. > 60% relative density), mortality exceeds recruitment and these forests exhibit self thinning (Cochran *et al.*, 1994).

The overall ESM density distribution closely approximated the -3/2 self thinning power law with a scaling exponent, *b* = -1.4±0.1 (Equation 1), using a 0.99-quantile regression. In ongoing work, we explore why this relationship remains contentious (Enquist & Niklas, 2001, West *et al.*, 2009). For example, at ~100% relative density, forests appeared to be best delineated by a more constrained thinning slope, *b* = -1.75 (Fig. 1a) using a linear 0.9 quantile regression through log-transformed self-thinning forests (i.e. relative density > 60%). Forests with the largest mean diameters (i.e. > 35 cm; 1.8% of the dataset) have slowed or reversed mean diameter change and carbon accumulation, as existing tree growth, resource limitation, disturbance, and recruitment were roughly balanced during the census interval.

Temperature, precipitation and soil moisture anomalies were associated with significant change (p < 0.05) in carbon accumulation, stem density and radial tree growth in eastern US forests (Fig. 2). Forests that experienced mean summer temperature anomalies +1°C (range: 0.8-2°C) above average over the 5-year census interval were associated with higher rates of disease (+23%, CI: 3%-42%), weather disturbance (+57%, CI: 27%-87%), and were >10% more likely to experience either an anomalously wet summer, dry summer or 5-year dry period (Tables S3 & S4). These warmer conditions appeared to have minimal influence on overall stem density and radial tree growth; but, forests experiencing these conditions had reduced mean diameter growth, lower sapling density, and a 27% reduction in productivity (−0.4±0.1 Mg C ha^-1^ yr^-1^, n = 759 plots). Excluding disturbances and saplings, warming had little effect on productivity (Fig. S1). Thus, forest productivity was reduced indirectly through enhanced disturbances and sapling mortality/lack of recruitment associated with warming conditions.

**Figure 2.**
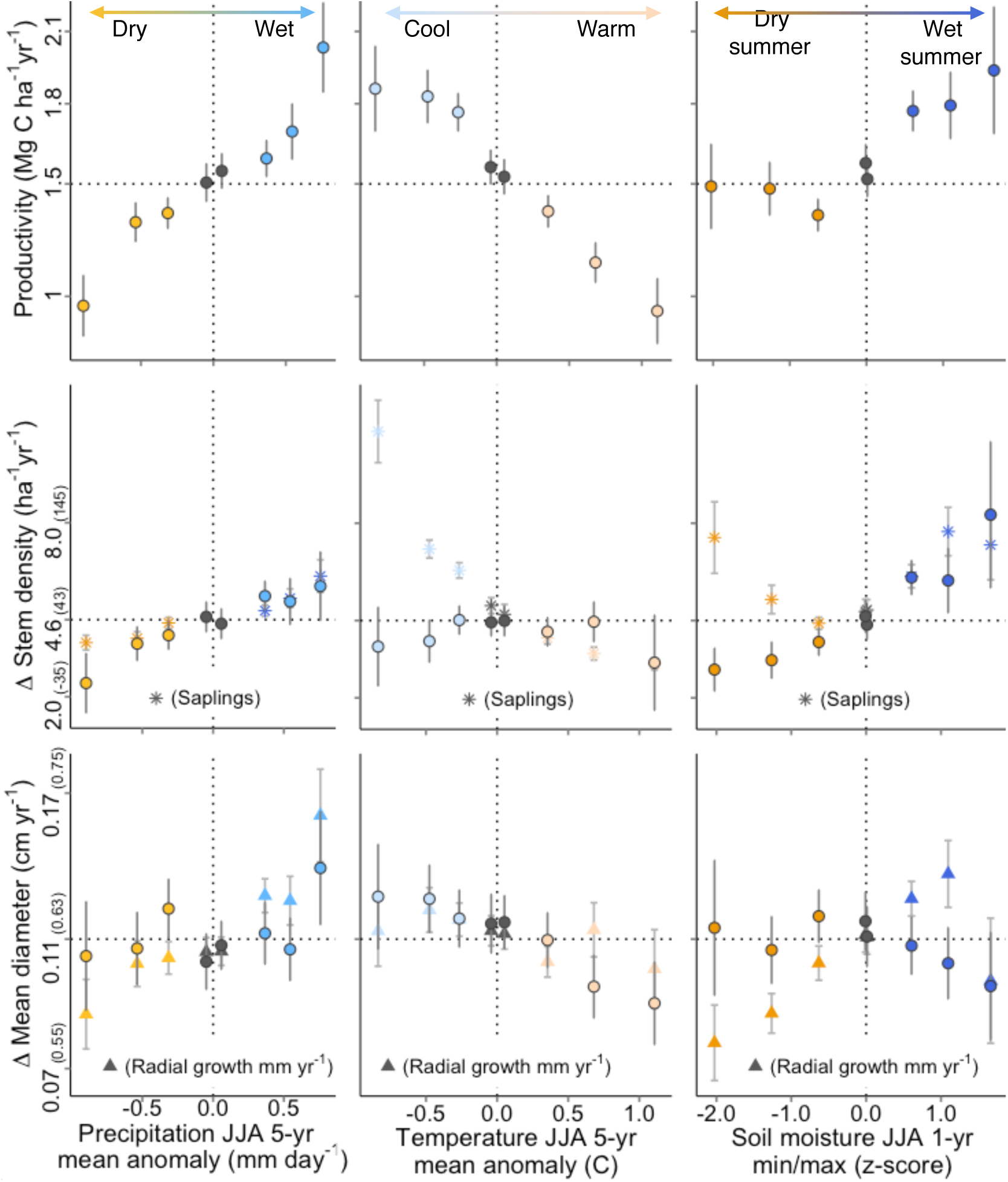
Influence of changing climate on forest structure. Change in productivity, stem density and radial tree growth in forests that experienced unusually wet, dry, warm or cool summer climatic conditions (June-August [JJA]) relative to ESM groupings that experienced non-anomalous conditions. Climate conditions were split into eight intervals (n > 100; AC scores: 0-2, 2-7, 7-17, 30-40 [grey], 60-70 [grey], 83-93, 93-98, 98-100). Points are independent with bootstrapped uncertainty (±1.96SE; 1000 iterations). Sapling stem density and radial growth of living trees are also shown with y-axis values in parentheses.

Cooling conditions (−0.8°C, range -0.6 to -1.7°C) were associated with an opposite, +30% increase in productivity (+0.4±0.2 Mg C ha^-1^ yr^-1^, n = 398 plots, Fig. 2) that was also partially driven by altered disturbance regimes and sapling dynamics (Fig. S1). The productivity changes associated with warming and cooling conditions were derived independent from each other. Five-year wet periods increased productivity by up to +34% (+0.6±0.1 Mg C ha^-1^ yr^-1^, n = 502) with +0.7 (SD: ±0.2) mm day^-1^ more summer precipitation; while, 5-year dry periods, decreased productivity by up to 34% (−0.5±0.1 Mg C ha^-1^ yr^-1^, n = 447) with 0.9 (SD: ±0.2) mm day^-1^ less summer precipitation. In contrast, 1-year dry summer conditions had a lasting influence on radial growth (measured over the 5-year census interval), but a weak influence on productivity, due in part because sapling stem density had the opposite response of larger trees. Stem density and radial growth were generally more sensitive to precipitation than warming. Deciduous and coniferous dominated forests had similar productivity relationships to climate conditions, however a sub-regional breakdown revealed warming effects were insignificant in the northern latitudes (45-49°N) where more frequent wet summers and 5-year wet periods may have offset any reduction in productivity (Figs. S2, S3, S4 & Table S4).

### Forward modeling

We used ESM as a basis for projecting current forest change into the future under different disturbance regimes and climate conditions. At the start of the future projections (Fig. 3a), representing approximately current conditions, the average carbon biomass of all FIA plots underestimates forest carbon sink potential due to the overabundance of plots recovering from land use change relative to those recovering from natural disturbances. As the model proceeds, the forward model selects forest plots that represent part of the natural density cycle (or with logging), from which it calculates the average mean diameter and carbon of the selected forests. Notably, because projections are empirically derived, they do not include future CO_2_ fertilization, however, as we discuss below, productivity enhancement may not control final carbon sink potential.

**Figure 3.**
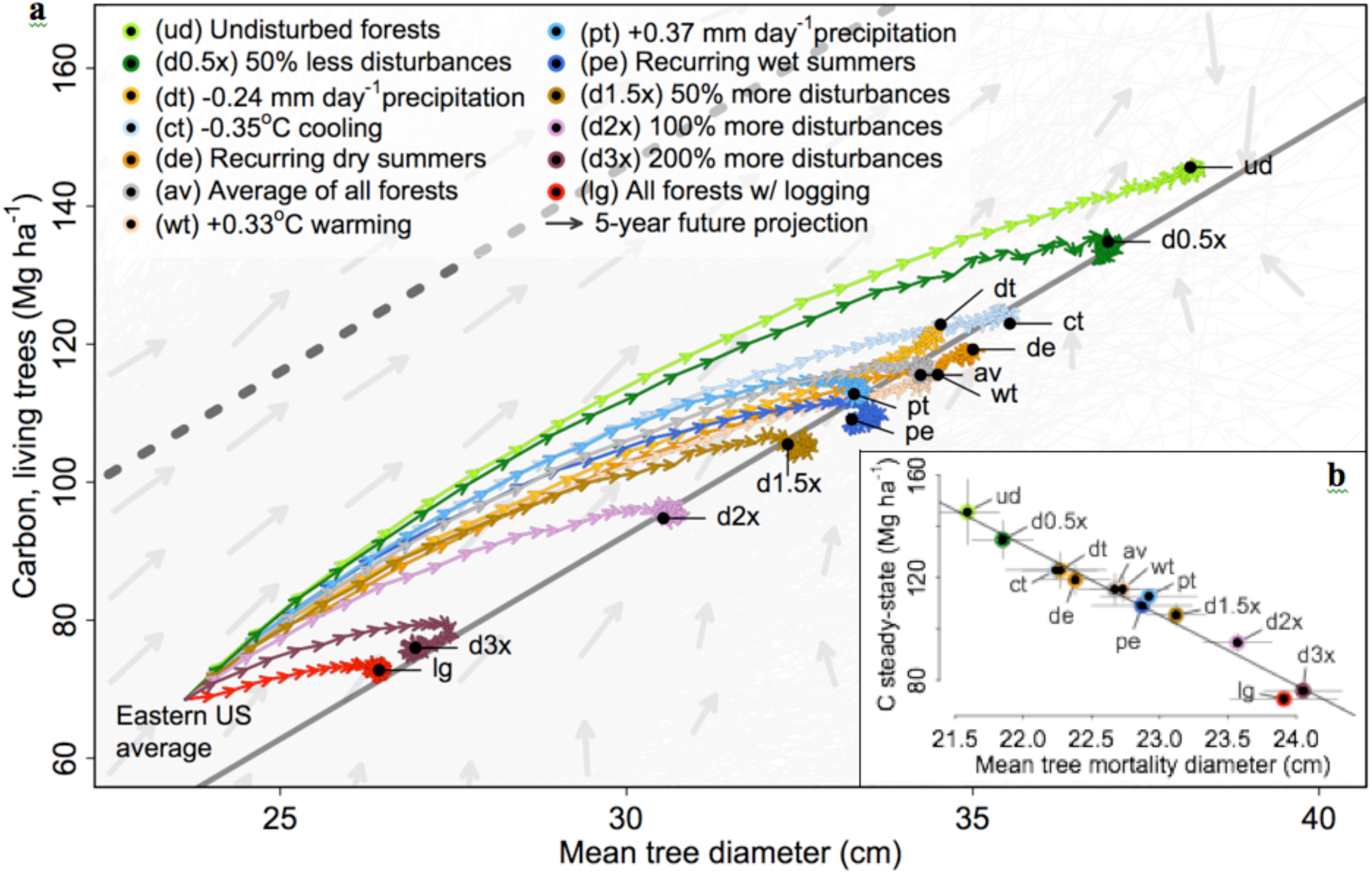
Carbon steady-state projections and relationship to tree mortality size. Eastern US temperate forest vegetation carbon was projected into the future at 5-year increments under moderate climate-change conditions and various disturbance intensities. Each scenario reached steady-state where density remains fairly constant after ~150 years. The average of all forests (av) excludes forests that were logged. For reference, species composition of the plots in the forward projections generally shifted toward mid- and late-successional species. The projected average eastern US forests mean tree diameter (D_plot_) and carbon steady-state were linearly related: C_sink_ = 5.9D_plot_ - 85; r^2^ = 0.98 (solid grey line). The projected C_sink_ was predicted by D_mortality_ following a linear relationship: C_sink_ = -27.3D_mortality_ + 734; r^2^ = 0.96 (b); shown with the standard error of mean mortality diameter (±1.96SE) and forward model uncertainty (±1.96SD; thin grey lines).

While steady-state vegetative carbon sink projections represent the range of filtered FIA sampled species/locations, the distribution of plot structures shifts to accommodate the structural implications of succession. For example, the ESM has more early successional forests than large disturbances that reset succession; and thus, through iterations of the forward model, the distribution of plots shifts to follow the cyclical pattern of the ESM (Fig. 1a), from recruitment through self-thinning and reduction in density followed by recruitment typical of gap dynamics in later successional forests. Late successional forests are subtly represented in the ESM density mapping where low density overstory recruitment and gap-infilling proceed toward mean diameter growth with less thinning at lower relative densities (expected for typical late successional forests) than typical mid-successional forests at maximum relative density. The difference between vegetative carbon sink projections that separate or mix mid and late successional species was minor as explained in Supplemental Information.

Under the breadth of conditions that forests experienced in the FIA dataset, average unmanaged forests of the eastern US would reach a steady-state density in ~145 years gaining +43 Mg C ha^-1^ for a maximum vegetation carbon sink (C sink potential) of 112±7 Mg C ha^-1^, with a density of 281±10 stems ha^-1^, mean tree diameter of 34±1 cm (Figs. 1 & 3). Management (i.e. logging), changing climate, and disturbance regimes produced substantial differences in future C sink potential ranging from 72±2 Mg C ha^-1^ with current eastern US logging regimes to 153±21 Mg C ha^-1^ in undisturbed forests (Fig. 3). C sink potential was linearly related to projected mean diameter (slope = 5.9), similar to the relationship found in Fig. 1b (slope = 5.6), which suggests C projections well represent the distribution of eastern US forests. These relationships represent the combination of allometric equations for forests of the eastern US (Jenkins *et al.*, 2003).

Surprisingly, C sink potential under higher precipitation conditions (wet) with enhanced productivity (i.e. Fig. 2) was below average, while warming and drying conditions had average to above average C sink potential. The projected growth of mean diameters and vegetative carbon sink potential were primarily related to the size of trees that died in thinning forests during the census interval (D_mortality_) in the underlying datasets (p < 0.001, r^2^ = 0.96; Fig. 3b). Simply, mortality of smaller trees (rather than larger trees) is significantly correlated to greater landscape-scale carbon storage in this model.

## DISCUSSION

The coherent, cyclic pattern of the ESM shows average structural changes become robust and predictable at broad spatial scales despite the immense complexity of forest habitats across eastern US. The ESM provides a new perspective on successional patterns, including early successional establishment and growth, self-thinning, and gap filling. The ESM was also useful as comparative tool to examine the relative influences of climate, disturbance and climate induced disturbance. Most striking was the strong negative influence of warming on forest productivity, and the significant correlation of warm anomalies to enhanced disturbances and sapling mortality/lack of recruitment. Results suggest near-term direct forest growth response to moderate warming will be limited compared to indirect responses through intensified disease, weather disturbances, and sapling dynamics in the eastern US overall. Notably, insect infestations were less common during both warming and drying conditions, contrary to expectations (Anderegg *et al.*, 2013) (Table S3), potentially because eastern US forest ecosystems are not generally moisture limited (Hanson *et al.*, 2001) despite radial growth being a sensitive indicator of drought and pluvials (Fig. 2; (Martin-Benito & Pederson, 2015). Further analyses on the potential for species sensitivities to this warming-disease/weather disturbance relationship could help inform forest management in a warming world.

This work adds to a growing body of literature that relates forest structural characteristics to patterns of change useful for predictive modeling. We expect the ESM framework could be expanded to consider of other forest characteristics and support other findings, such as relationships to wood density (Woodall *et al.*, 2015), Stand Density Index (Woodall *et al.*, 2005), species composition and demography (Vanderwel *et al.*, 2017), or plant functional types (proposed as future work). Our findings indicate the complex role of climate as a driver of forest change (e.g. Lichstein *et al.*, 2014) may need to further consider the relationships between climate and forest disturbance.

### Forward modeling

Future scenarios of forest carbon sequestration can inform climate mitigation planning and forest management, particularly in the context of changing climate (Caspersen *et al.*, 2000). Surprisingly, the influence of climate on productivity was not directly related to future C sink potential, for example higher precipitation and higher soil moisture conditions had enhanced productivity (i.e. Fig. 2) but below average C sink potential. Instead the results suggest that when forests are resource limited, conditions that induce mortality of smaller trees (rather than larger trees) lead to greater landscape-scale carbon storage during the ~150 year projection of this study (although longer-term transitions to late succesional species could be impacted). While we know large trees uptake more carbon (Stephenson *et al.*, 2014), the relationship applied to scenarios that both enhanced forest productivity (e.g. wet summers) and reduced forest productivity (e.g. 5-yr dry periods), which suggests mortality-size influences long-term carbon storage in living biomass more than factors that alter productivity (e.g. soil moisture, CO_2_ fertilization).

There are a number of forward modeling limitations. First, the carbon steady-state projections and the relationships to tree mortality size represent our best estimate with uncertainty for the projected ~150 years. Mid-successional forests were well sampled by the FIA and future vegetative carbon sink projections were generally within the lifespan of these species. In the very distant future, the carbon steady-state may continue to shift as forests shift toward late-successional species, depending on, in part, disturbance regimes (Thom *et al.*, 2017). Second, while the future of long term climate change remains difficult to predict, one possible situation is for climate anomalies to continue becoming more and more extreme. Thus the anomalies used in this research that are of short duration and particularly unusual over the last century (e.g. many of the warmest years on recorded occurred since 1998 globally) may become more and more common and intense. Alternatively, the forward modeling of repeated anomalies may overestimate impacts. Nonetheless, the primary value of the scenarios presented are to show a range of conditions that could feasibly occur, given model limitations, and highlight the need for more research to understand factors that influence average tree mortality size.

Because the moderate climate change scenarios and the more extreme disturbance scenarios were linearly related (Fig. 3a), we suggest that changing disturbance characteristics were a common mechanism determining C sink potential. Interestingly, the disturbance characteristics of the warming scenario (+0.33°C) did not shift mean mortality-size or C sink potential. However for reference, greater warming (+1°C) was associated with mortality of larger trees (+1 cm mean mortality size, compared to non-AC forests). Thus, following the relationship to mean mortality size found in Fig. 3, climate warming of +1°C could enhance disturbance of larger trees and indirectly reduce temperate forest future C sink potential by up to ~15%. Much work is needed to further understand factors driving change in mortality-size and how these relationships may change in the future. In particular, improved disturbance prediction is critically important to narrow the range of C sink potential we found (i.e. 72 - 153 Mg C ha^-1^).

The most extreme disturbance scenario (3x modern disturbance rates) had a similar C sink potential to the actively managed forest scenario, which suggests that efforts to manage for resilience (Dale *et al.*, 2001) could effectively establish the minimum average C sink potential for the vegetative landscape. Thus active management (e.g. thinning, salvage logging) can drastically reduce the uncertainty of carbon storage in landscapes. The future of unmanaged forests (e.g. many national parks), is less certain and we expect a warming climate will continue to indirectly impact temperate forests by enhancing mortality attributable to extreme weather and disease (Fig. 2). Overall, this work intends to describe general patterns of large-scale changes. More refined approaches are needed, for example, to understand the interaction of climate variability and other factors that drive non-linear disturbance spread (i.e. mega-wildfires).

## Acknowledgments

We acknowledge the immense field and data management efforts provided by FIA staff that made these analyses possible. Thanks to Ben Felzer, the Walt Cason lab, and numerous US Forest Service staff in the Northeastern Area office, for early review. MCD supported by NSF Macrosystems 1318164.

## Author Contributions

T.D.A., M.C.D. and R.K.B designed the project. T.D.A. performed analyses and wrote the initial paper. All authors revised and commented on the manuscript.

## Author Information

FIA data are publicly available from apps.fs.fed.us/fiadb-downloads/datamart.html. Climate data provided by the NOAA/OAR/ESRL PSD, Boulder, Colorado, USA, from their website at http://www.esrl.noaa.gov/psd/. Analysis R code is accessible via Github (github.com/wanderswest/ESM-FIA). The authors declare no competing financial interests. Correspondence and requests for materials should be addressed to.

## Supplementary Information

### ESM Climate sensitivity

The ESM groupings were created as a basis for comparison for density, mean diameter, radial growth, and carbon change under specific climate conditions. Generally, comparisons of structural trajectories were robust across temperate forest types and mean temperatures, however, the magnitude of structural change was sensitive to these factors. Therefore data used to build the comparison non-anomalous climate (non-AC) ESM required a few additional considerations to maximize sensitivity and minimize biases, described below. We expect climate anomalies had a consistent effect across microclimates.

Data for the non-AC ESM was selected from the same 0.5°x0.5° grid cells with the same weighted frequency of sampling as in the AC data. This spatial weighting was based on a simple weighted average using the proportion of AC forest plots to non-AC forest plots within each 0.5°x0.5° grid cell; technically defined below:

#### Non-AC ESM spatial weighting

- Grid cell proportion = Number of AC plots / total number of plots; within each latitude/ longitude grid cell
- ESM grouping proportional mean = mean(Grid cell proportion); within each ESM grouping
- Within each ESM grouping: (median climate plot structural change * Grid cell proportion)/ESM group proportional mean
- Structural changes of AC plots were then compared to this spatially weighted ESM.

Note, this simple weighting does not require plots in each grid cell to represent all ESM groupings and anomalous climate conditions (which would be ideal); but rather the weighting shifts the overall distribution of plots sampled, such that, any residual mismatch between these factors would be random and contribute toward analysis error. For reference, analyses were also run without spatial weighting, which produced similar results with larger uncertainty (results not provided). Additionally, a separate warming climate sensitivity analysis was also run comparing change in productivity at each AC plot to change in the three closest non-AC plots (in distance) that were of similar mean diameter, relative density and forest type (deciduous or coniferous). Results of this alternative analysis (not provided) showed reductions in productivity associated with warming that were within error of the provided results.

The independent intervals of climate conditions used in the climate anomaly analyses presented in Fig. 2 were chosen to clearly represent relationships. More/smaller intervals and moving-average intervals were also tested and produced similar results with greater variability or complexity. Rates of change were calculated per year by dividing change by the census interval. Bootstrapped confidence intervals (1.96SE, 1000 iterations) were applied to the difference between the trajectories of all the AC data from the associated non-AC ESM. AC datasets were smaller than non-AC ESMs with larger standard error, and as such, significance was determined if AC data confidence intervals did not overlap the non-AC model mean represented by the horizontal dashed line in Fig. 2 (i.e. results were significantly different from zero).

Climate conditions are expected to influence forest change through direct mechanisms (e.g. Q10 relationships) and by correlating with disturbance and other climate conditions (e.g. drought during warm periods). In Supplementary Table 3, for each climate condition, the likelihood of also experiencing specific disturbances relative to the likelihood of occurrence in the non-AC dataset is given (“Difference (%)”) with the number of occurrences (“n”). Here the ESM was not used as the basis for comparison. This comparison simply identifies if forests that experienced anomalous climate had significantly different frequency of mortality associated with disturbance agents than non-AC forests with the same spatial distribution (using a 0.5°x0.5° grid). Confidence intervals for the “Difference (%)” were determined by bootstrapping (1000 iterations) and disturbance analyses were repeated excluding forests with unknown disturbances to validate this large source of ambiguity had little influence on changes in the relative frequency of known disturbances (i.e. the results were generally similar). This same analysis was also performed to compare the relationship between AC conditions and other AC conditions (Supplementary Table 4).

### Forward steady-state model additional rational

Successional stages provide a well-known context for our findings but are not critical to delineate within the ESM for analyses. Structurally, size-density scaling relationships should broadly apply independent of tree size distribution and successional status (Enquist & Niklas, 2001). Future projections using the ESM were appropriate for four reasons: 1) The FIA samples a comprehensive range of size-density relationships including even-aged and uneven-aged forests. Saplings (i.e. DBH < 12.7 cm) were excluded from the ESM density mapping to substantially reduce the resolution of forest structural complexity, such that mean arithmetic diameter, mean geometric diameter, and quadratic mean diameter all produce highly similar versions of the ESM. The exclusion of saplings produced an ESM where forests with the greatest biomass occupied the largest mean diameters (rather than being biased toward the center of the mean diameter range if small saplings were included). 2) Relative density (stocking) calculations were performed by the FIA based, in part, on standards to maximize timber production and thus fully- and over-stocked forests would generally be early and mid successional forests and late successional forest will generally occupy lower relative densities. 3) Where mid and late successional type forests overlap in ESM mean diameter/relative density groupings, the forward model would select either type indiscriminately. For reference, in species-level analyses (not shown) we found forests dominated by mid successional species, have larger mean diameters but similar carbon content as compared to forests dominated by late successional species with similar relative densities (i.e. within an ESM grouping, mid successional species would tend to have larger mean diameters than late successional species but the change in carbon content would be similar). Thus any mix between mid and late successional species in the forward model may overestimate equilibrium mean diameters but should remain an accurate predictor of forest carbon sink potential.

**Supplementary Figure 1.**
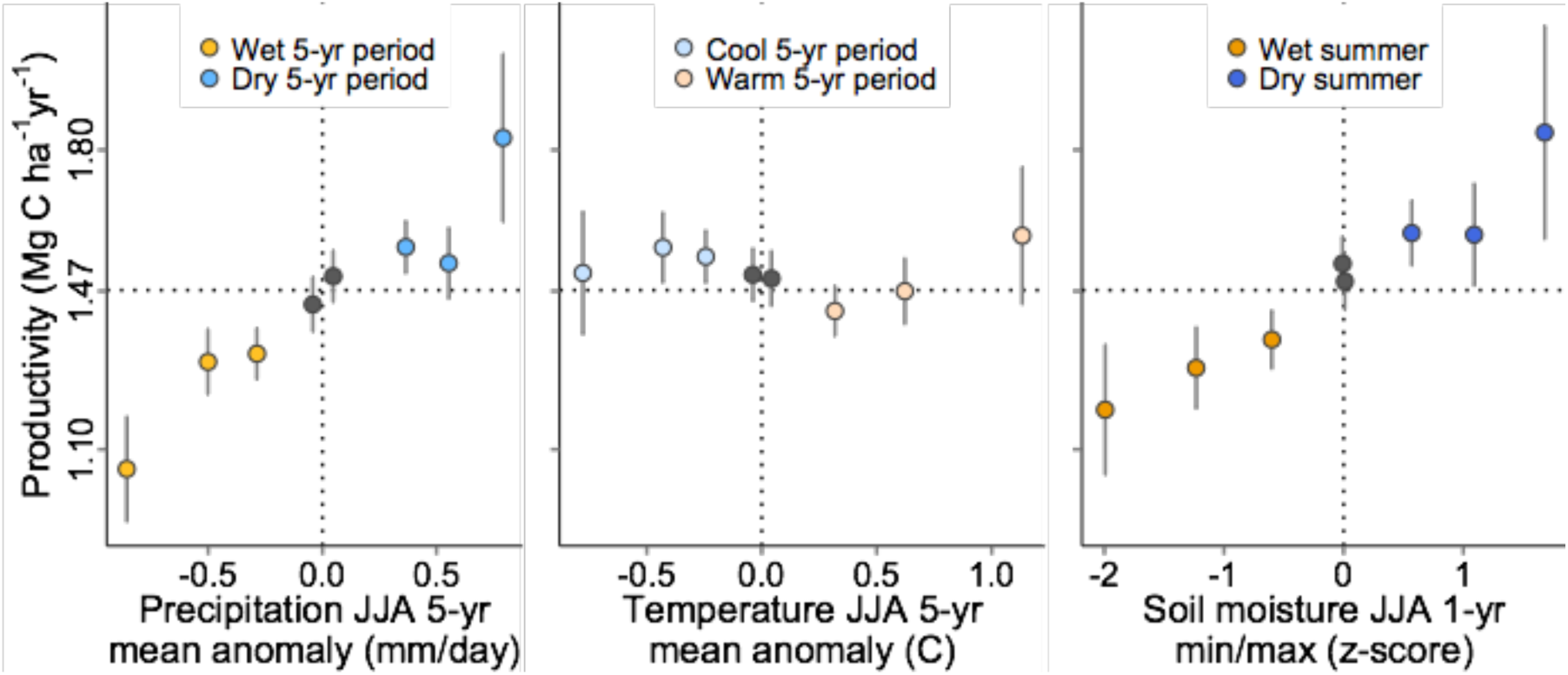
Change in productivity for forests without disturbances and excluding saplings for climate conditions relative to a comparison non-AC ESM (represented by the horizontal dotted line). Points are independent averages (n > 100) at intervals of climate divergence (AC scores: 0-5, 5-12.5, 12.5-25, 30-40, 60-70, 75-87.5, 87.5-95, 95-100) with bootstrapped uncertainty (±1.96SE; 1000 iterations).

**Supplementary Figure 2.**
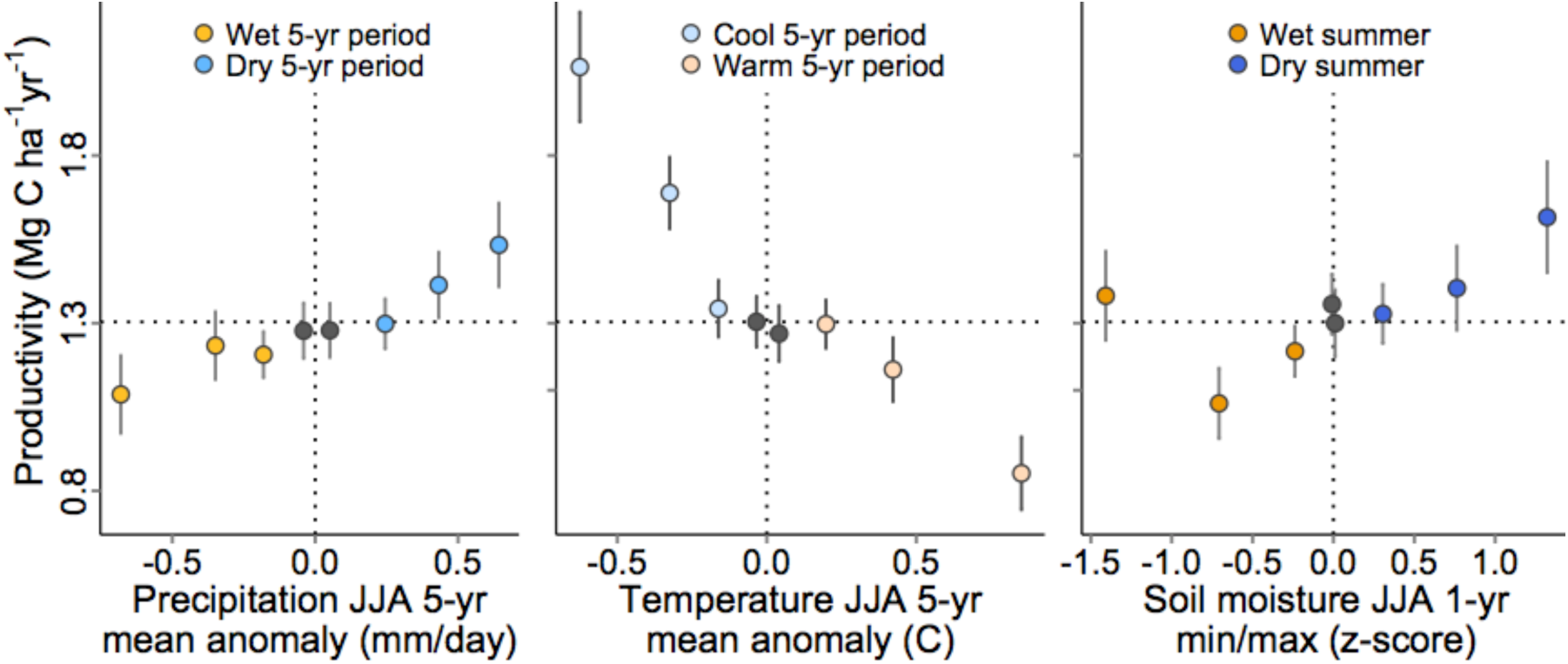
Deciduous forest change in productivity for climate conditions relative to a comparison non-AC ESM (represented by the horizontal dotted lines). Points are independent averages (n > 100) at intervals of climate divergence (AC scores: 0-5, 5-12.5, 12.5-25, 30-40, 60-70, 75-87.5, 87.5-95, 95-100) with bootstrapped uncertainty (±1.96SE; 1000 iterations).

**Supplementary Figure 3.**
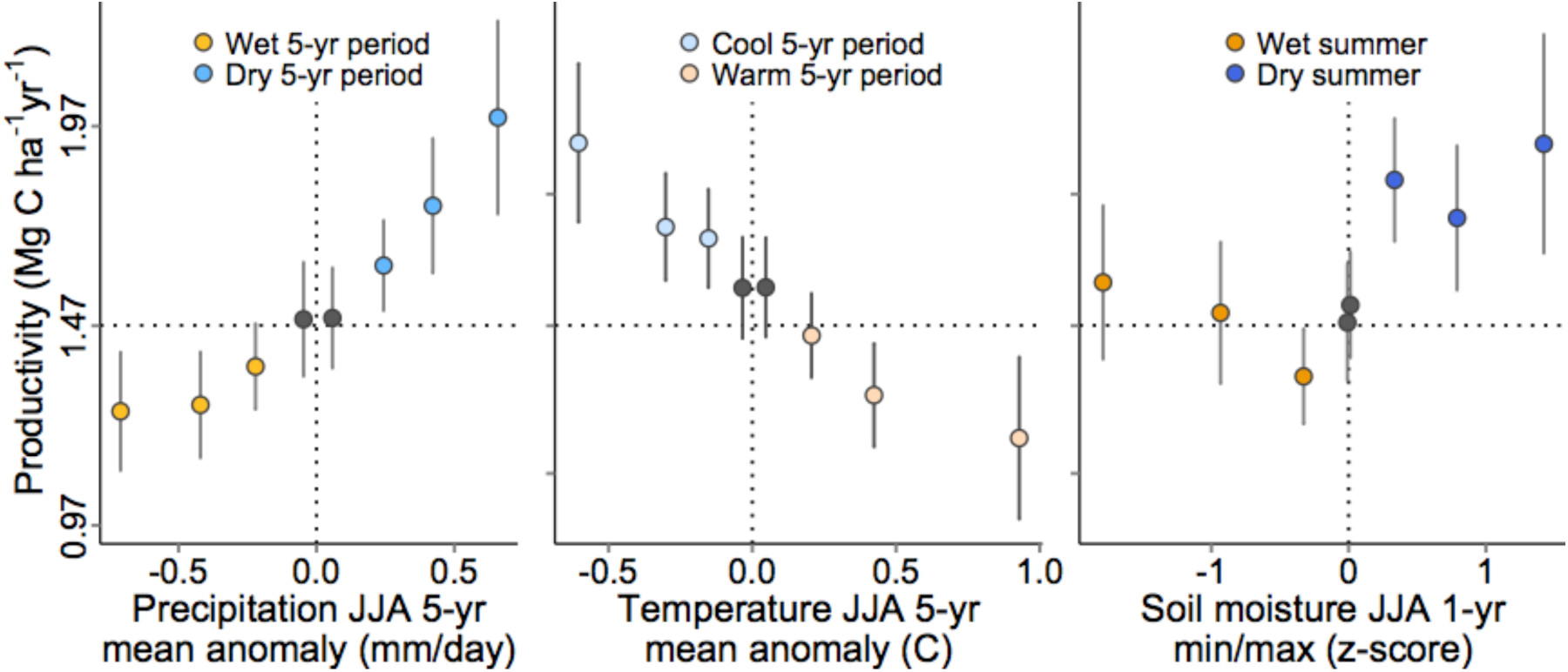
Coniferous forest change in productivity for climate conditions relative to a comparison non-AC ESM (represented by the horizontal dotted line). Points are independent averages (n > 100) at intervals of climate divergence (AC scores: 0-5, 5-12.5, 12.5-25, 30-40, 60-70, 75-87.5, 87.5-95, 95-100) with bootstrapped uncertainty (±1.96SE; 1000 iterations). Notably, a dominant forest species was not attributed by the FIA to 15% of the dataset.

**Supplementary Figure 4.**
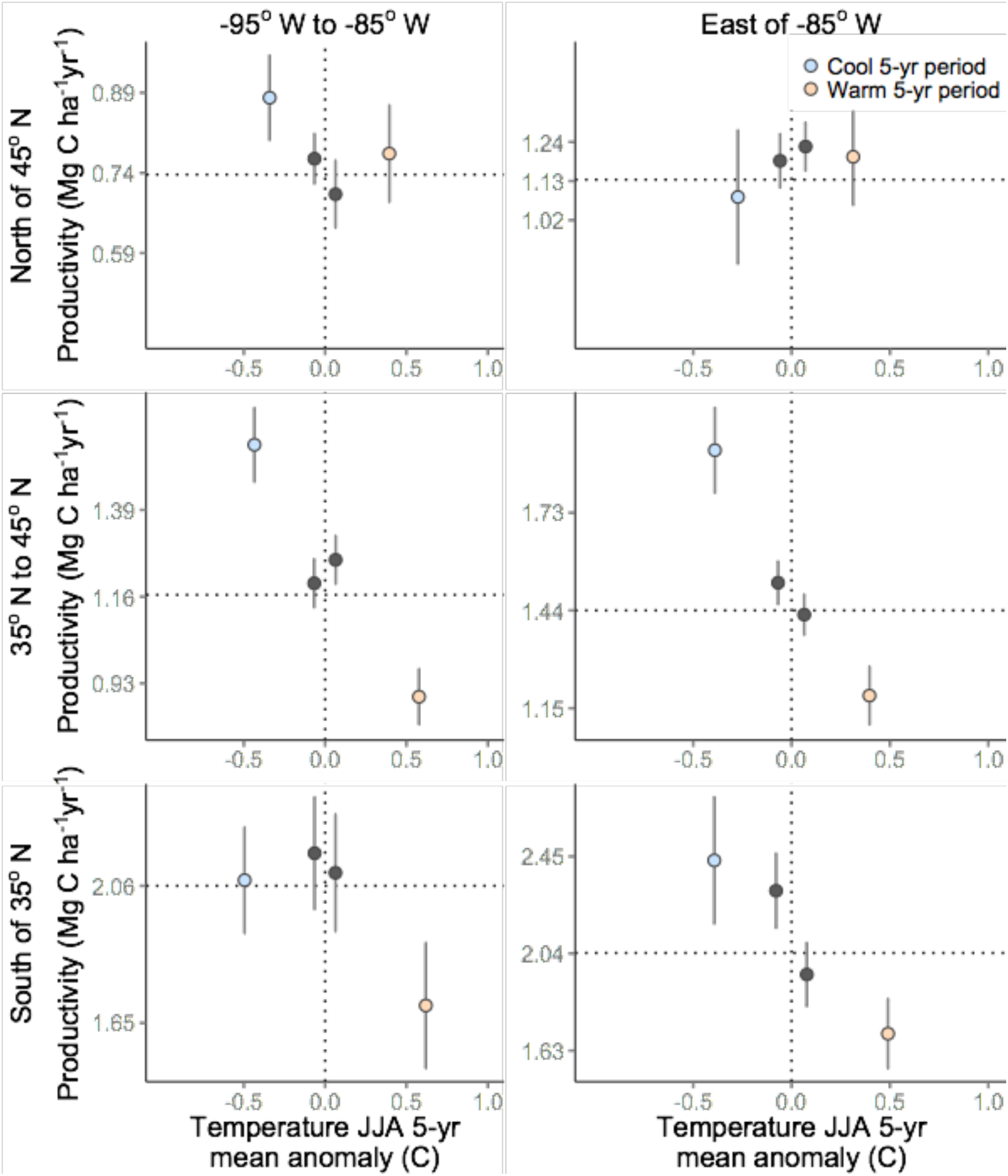
Sub-regional analysis of temperature anomalies and productivity. Points are independent averages (n > 100) at absolute intervals of climate divergence: less than -0.2°C, between -0.2°C and 0°C, between 0°C and 0.2°C, and greater than 0.2°C; with bootstrapped uncertainty (±1.96SE; 1000 iterations). North of 45°N, warming impacts may have been offset by an extended growing season or correlation with increased precipitation (see table S4).

**Supplementary Table 1.** FIA data filters used to select forest plots for density change analysis.

| Data filter goal | FIA filter method | FIA/R reference |
| --- | --- | --- |
| Sample forest interiors and exclude sharp edge ecotones | Plots have only 1 condition | CONDmax = 1 |
| Plots have tree(s) that have been resurveyed | Plots had at least one tree at first survey | PREV_STOCKING > 0 |
| Exclude plots with previous survey error(s) | Exclude plots with reconciled trees i.e. RECONCIDCD>4 | STATUSCD ≠ 0 or 3 |
| Exclude plots with trees unknown to be alive or dead | All trees must have a status | STATUSCD = 1 or 2 |
| Plots are all the same size | Plots are all national standard FIA design | DESIGNCD = 1 |
| Plots resurveyed after ~5 years and outliers removed | Exclude plots resurveyed over less than 3 years or more than 9.5 years | REMPER ≥ 3 and ≤ 9.5 |
| Exclude saplings and seedlings | Exclude trees with DIA<5in. (12.3 cm) | DIA ≤ 5 |
| Eastern US defined as east of -95 W longitude | Exclude trees west of -95W | LON < -95 |
| Only plots used by USFS to calculate growth | Merge subset data with all live trees from the TREE_GRM_ESTN table | ESTN_TYPE = AL |
| Use only breast height tree diameter measurements | Use DIA measurements from TREE_GRM_ESTN table | DIA = DBHcalc |
| Exclude artificially regenerated forests | Select only naturally occurring forests | STDORGCD = 0 |
| Exclude trees not part of the resurvey plot | Exclude any trees not used in the TREE_GRM_ESTN table | P2A_GRM_FLG ≠ "N" |

**Supplementary Table 2.** Climate anomaly descriptions, mean condition values and sample size (n). Most AC conditions = AC scores of 0-3 or 97-100. Moderate AC conditions = AC scores 3-17 or 83-97, see Table 1 for scoring methodology.

| Climate condition | Description | Steady-state model (Ss) | Moderate AC | Most AC |
| --- | --- | --- | --- | --- |
| Wet 5-year period | Anomalously high mean June-August precipitation during the five-years prior to forest resurvey. | 0.37 mm/day<br>n = 9979 | 0.44 mm/day<br>n = 4970 | 0.68 mm/day<br>n = 513 |
| Wet summer | Maximum z-scored mean June-August soil moisture that occurred between 1-5 years prior to forest resurvey. | 0.69 (z-score)<br>n = 7582 | 0.8 (z-score)<br>n = 4110 | 1.32 (z-score)<br>n = 750 |
| Dry 5-year period | Anomalously low mean June-August precipitation during the five-years prior to forest resurvey. | -0.24 mm/day<br>n = 15484 | -0.39 mm/day<br>n = 5541 | -0.61 mm/day<br>n = 985 |
| Dry summer | Minimum z-scored mean June-August soil moisture that occurred between 1-5 years prior to forest resurvey. | -0.53 (z-score)<br>n = 14450 | -0.78 (z-score)<br>n = 4568 | -1.37 (z-score)<br>n = 1481 |
| Warm 5-year period | Anomalously high mean June-August temperature during the five-years prior to forest resurvey. | +0.33°C<br>n = 10308 | +0.50°C<br>n = 4945 | +1.0°C<br>n = 759 |
| Cool 5-year period | Anomalously low mean June-August temperature during the five-years prior to forest resurvey. | -0.35°C<br>n = 8593 | -0.36°C<br>n = 5023 | -0.62°C<br>n = 1161 |

**Supplementary Table 3.**
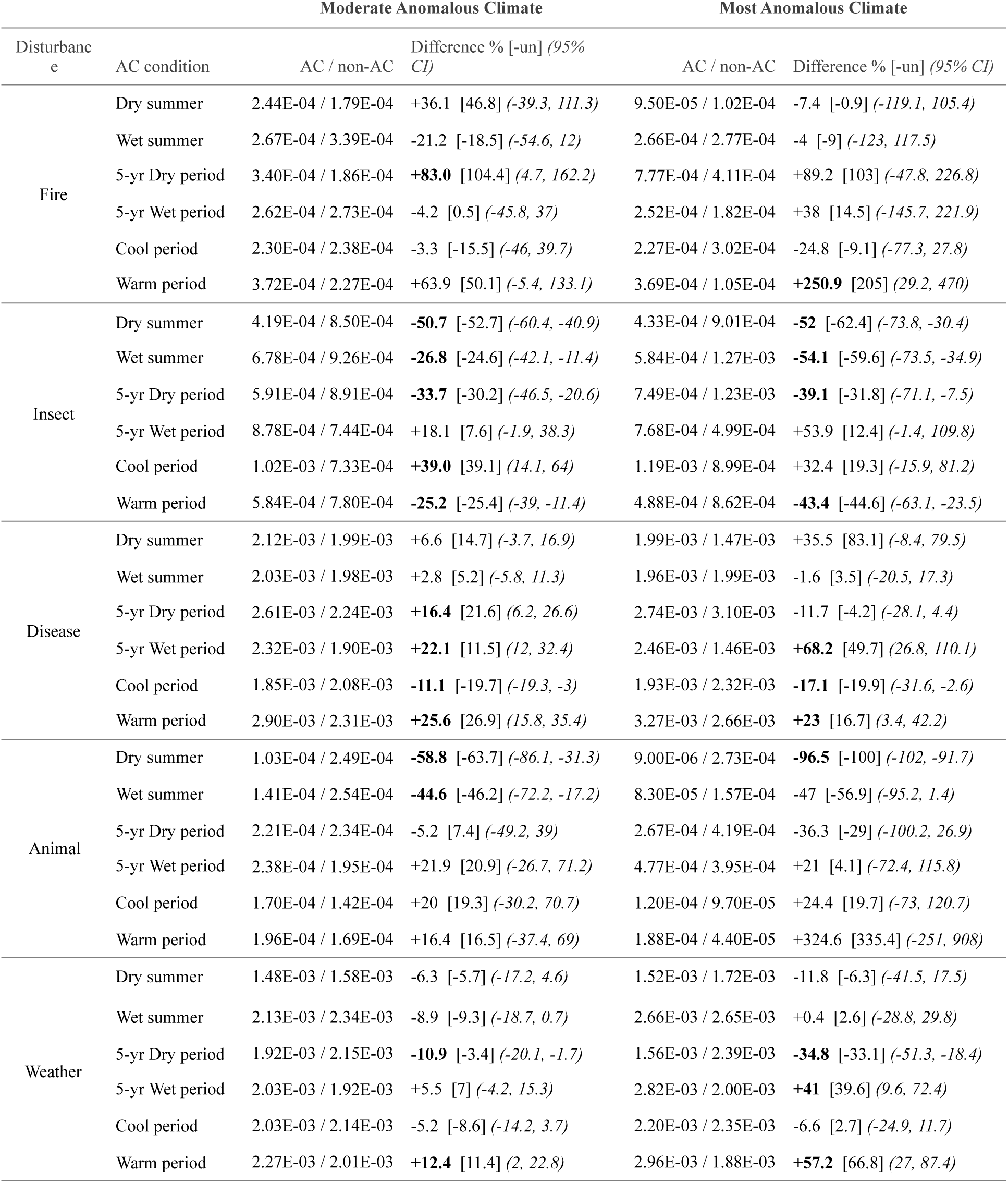
Correlation of disturbances with climate conditions using the percent difference between the mean proportion of disturbed/undisturbed trees on plots under AC and non-AC conditions (AC / non-AC). The AC conditions were further defined in Supplementary Table 2. Bold values indicate significance (p < 0.05) based on bootstrapped 95% confidence intervals (10,000 iterations). Percent difference calculations were repeated excluding forests with unknown disturbances [-un]. For example, the last line of the table indicates in the same geographical area and census interval, the ratio of plots that experienced mortality attributed to weather (e.g. wind storm) and a moderately warm period (AC score 83-97) was 0.00227, whereas the ratio of those with weather disturbance and without warming was 0.00201, thus suggesting weather disturbances were 12.4% more likely to occur during the same census interval as warm periods with a 95% confidence interval from 2% - 22.8%.

**Supplementary Table 4.**
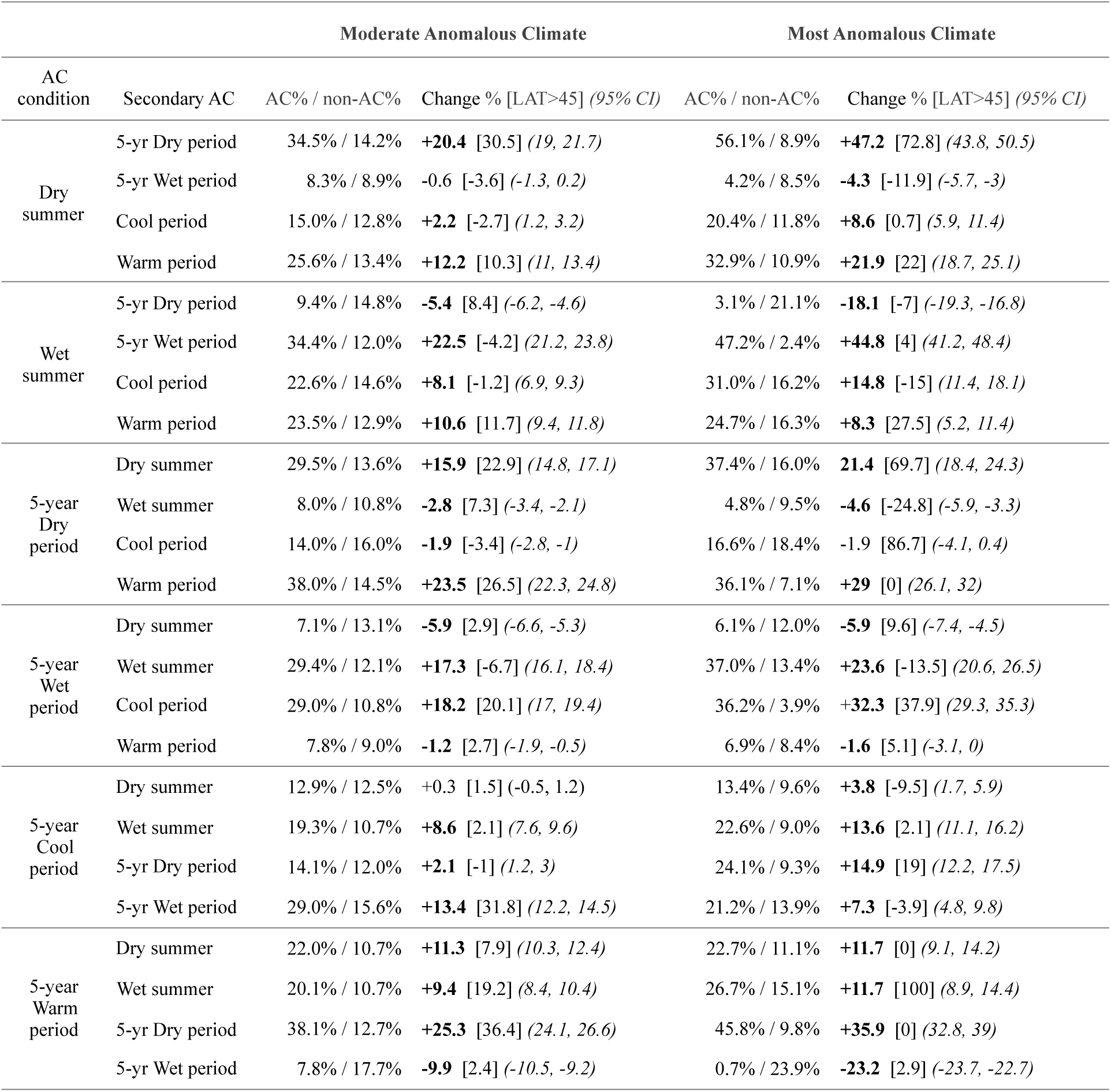
The frequency of Anomalous Climate (AC) occurring during the same census interval as other AC climate conditions (Secondary AC = Moderate AC or Most AC). Bootstrapped 95% confidence intervals (10,000 iterations) of the difference calculation were provided and bold values indicate significance (p < 0.05). Percent difference calculations were repeated for eastern US forests above 45°N latitude [LAT>45]. For example, the first line of the table indicates in the same geographical area and census interval, 34.5% of plots experienced both a Moderate AC dry summer and a 5-year dry period, whereas without a dry summer (AC scores 30-70), only 14.2% of plots experienced a 5-year dry period; thus forests that experienced a moderately dry summer were 20.4% more likely to also experience a 5-year dry period. Similarly, forests that experienced a Most AC dry summer (AC scores 0-3), were 47.2% more likely to also experience a 5-year dry period [and 72.8% more likely north of 45°N latitude].

| AC condition | Moderate Anomalous Climate |  |  | Most Anomalous Climate |  |
| --- | --- | --- | --- | --- | --- |
|  | Secondary AC | AC% / non-AC% | Change % [LAT>45] (95% CI) | AC% / non-AC% | Change % [LAT>45] (95% CI) |
| Dry summer | 5-yr Dry period | 34.5% / 14.2% | **+20.4** [30.5] (19, 21.7) | 56.1% / 8.9% | **+47.2** [72.8] (43.8, 50.5) |
|  | 5-yr Wet period | 8.3% / 8.9% | -0.6 [-3.6] (-1.3, 0.2) | 4.2% / 8.5% | **-4.3** [-11.9] (-5.7, -3) |
|  | Cool period | 15.0% / 12.8% | **+2.2** [-2.7] (1.2, 3.2) | 20.4% / 11.8% | **+8.6** [0.7] (5.9, 11.4) |
|  | Warm period | 25.6% / 13.4% | **+12.2** [10.3] (11, 13.4) | 32.9% / 10.9% | **+21.9** [22] (18.7, 25.1) |
| Wet summer | 5-yr Dry period | 9.4% / 14.8% | **-5.4** [8.4] (-6.2, -4.6) | 3.1% / 21.1% | **-18.1** [-7] (-19.3, -16.8) |
|  | 5-yr Wet period | 34.4% / 12.0% | **+22.5** [-4.2] (21.2, 23.8) | 47.2% / 2.4% | **+44.8** [4] (41.2, 48.4) |
|  | Cool period | 22.6% / 14.6% | **+8.1** [-1.2] (6.9, 9.3) | 31.0% / 16.2% | **+14.8** [-15] (11.4, 18.1) |
|  | Warm period | 23.5% / 12.9% | **+10.6** [11.7] (9.4, 11.8) | 24.7% / 16.3% | **+8.3** [27.5] (5.2, 11.4) |
| 5-year Dry period | Dry summer | 29.5% / 13.6% | **+15.9** [22.9] (14.8, 17.1) | 37.4% / 16.0% | **21.4** [69.7] (18.4, 24.3) |
|  | Wet summer | 8.0% / 10.8% | **-2.8** [7.3] (-3.4, -2.1) | 4.8% / 9.5% | **-4.6** [-24.8] (-5.9, -3.3) |
|  | Cool period | 14.0% / 16.0% | **-1.9** [-3.4] (-2.8, -1) | 16.6% / 18.4% | -1.9 [86.7] (-4.1, 0.4) |
|  | Warm period | 38.0% / 14.5% | **+23.5** [26.5] (22.3, 24.8) | 36.1% / 7.1% | **+29** [0] (26.1, 32) |
| 5-year Wet period | Dry summer | 7.1% / 13.1% | **-5.9** [2.9] (-6.6, -5.3) | 6.1% / 12.0% | **-5.9** [9.6] (-7.4, -4.5) |
|  | Wet summer | 29.4% / 12.1% | **+17.3** [-6.7] (16.1, 18.4) | 37.0% / 13.4% | **+23.6** [-13.5] (20.6, 26.5) |
|  | Cool period | 29.0% / 10.8% | **+18.2** [20.1] (17, 19.4) | 36.2% / 3.9% | **+32.3** [37.9] (29.3, 35.3) |
|  | Warm period | 7.8% / 9.0% | **-1.2** [2.7] (-1.9, -0.5) | 6.9% / 8.4% | **-1.6** [5.1] (-3.1, 0) |
| 5-year Cool period | Dry summer | 12.9% / 12.5% | +0.3 [1.5] (-0.5, 1.2) | 13.4% / 9.6% | **+3.8** [-9.5] (1.7, 5.9) |
|  | Wet summer | 19.3% / 10.7% | **+8.6** [2.1] (7.6, 9.6) | 22.6% / 9.0% | **+13.6** [2.1] (11.1, 16.2) |
|  | 5-yr Dry period | 14.1% / 12.0% | **+2.1** [-1] (1.2, 3) | 24.1% / 9.3% | **+14.9** [19] (12.2, 17.5) |
|  | 5-yr Wet period | 29.0% / 15.6% | **+13.4** [31.8] (12.2, 14.5) | 21.2% / 13.9% | **+7.3** [-3.9] (4.8, 9.8) |
| 5-year Warm period | Dry summer | 22.0% / 10.7% | **+11.3** [7.9] (10.3, 12.4) | 22.7% / 11.1% | **+11.7** [0] (9.1, 14.2) |
|  | Wet summer | 20.1% / 10.7% | **+9.4** [19.2] (8.4, 10.4) | 26.7% / 15.1% | **+11.7** [100] (8.9, 14.4) |
|  | 5-yr Dry period | 38.1% / 12.7% | **+25.3** [36.4] (24.1, 26.6) | 45.8% / 9.8% | **+35.9** [0] (32.8, 39) |
|  | 5-yr Wet period | 7.8% / 17.7% | **-9.9** [2.4] (-10.5, -9.2) | 0.7% / 23.9% | **-23.2** [2.9] (-23.7, -22.7) |

